# Oxygen-generating cryogel vaccines help overcome tumor antigen tolerance and induce durable anti-tumor immunity in prostate cancer

**DOI:** 10.64898/2026.05.13.724871

**Authors:** Alexandra Nukovic, Khushbu Bhatt, Thibault Colombani, Emilia Todorovic, Lucy M. Williamson, Brittany Noonan, Enrique M. Chang, Laura Losada Miguens, Michail Sitkovsky, Sidi A. Bencherif, Stephen M. Hatfield

## Abstract

Therapeutic cancer vaccines represent a promising approach to boost patients’ own immune system to fight cancer. However, many vaccine candidates have shown limited success in clinical trials in large part due to the insufficient antigen delivery to overcome tolerance and hypoxia mediated immunosuppressive mechanisms. Cryogel-based delivery scaffolds have emerged as a promising platform for cancer vaccines due to their biocompatibility and macroporous structure that allows for effective delivery to infiltrating antigen-presenting cells. However, these systems are limited by rapid, diffusion-mediated burst release of encapsulated recombinant proteins and local hypoxia-driven immunosuppression within the scaffold. Herein, we demonstrate that click conjugation of a tumor-associated protein within cryogel-based vaccines, combined with our new O_2_-generating platform (Click O_2_-Cryogel_VAX_), helps overcome immune suppression and weak antigenicity and primes effective anti-cancer immune responses. Sustained antigen delivery promotes cellular memory and Th1-mediated anti-cancer responses. By reversing hypoxia-driven immunosuppression, O_2_ acts as a powerful co-adjuvant to enhance humoral immunity. Together, Click O_2_-Cryogel_VAX_ supports a robust antitumor response that inhibits tumor growth and prolongs survival in a therapeutic prostate cancer model. These findings support the further research and development of Click O_2_-Cryogel_VAX_ as an effective delivery platform for therapeutic cancer vaccines.

## Introduction

Renewed interest in therapeutic cancer vaccines has followed major developments in vaccine technology and breakthroughs in cancer immunotherapy including immune checkpoint inhibitors (ICIs) and adoptive cell therapy (ACT).^1,2^ Prostate cancer is one of the leading causes of cancer-associated death in men, with a 5-year survival rate of 30 % in patients with metastatic castration-resistant prostate cancer (mCRPC).^3–5^ In response, several therapeutic vaccines targeting prostate cancer have been developed to help prime an immune response to reject recurrent disease or delay progression.^4,6^ Sipuleucel-T (PROVENGE^®^), an antigen-loaded dendritic cell (DC)-based vaccine, was the first official cancer vaccine/cell-based immunotherapy approved by the Food and Drug Administration (FDA) in 2010 but has had limited success.^7–10^ ProstVac-VF, a viral-based recombinant vaccine, showed improved median overall survival and was easier to administer as an “off-the-shelf” therapy compared to Sipuleucel-T; however, it showed no improvement as a monotherapy in Phase III trials.^8,9,11,12^ Recent advances in mRNA vaccine technology have also led to the exploration of neoadjuvant and personalized therapeutic cancer mRNA vaccines.^13,14^ BioNTech recently completed a Phase I/IIa trial investigating BNT112, a mRNA lipoplex vaccine as a monotherapy and in combination with anti-PD-1 (NCT04382898), while this trial was promising, Phase III trials will be required to reveal the clinical efficacy of the platform.^15^ Despite their ability to robustly boost B-cell and T-cell immunity against tumor cells, therapeutic cancer vaccines are limited by inadequate delivery methods, tumor heterogeneity, and the immunosuppressive tumor microenvironment (TME).^2,6,13^ It is clear now that success will require not only better formulations of antigen delivery vehicles but also innovative strategies to overcome tolerance and immunosuppression.^16–19^

To overcome immunosuppression and immune tolerance, and maintain effective humoral and cellular immunity against tumor-associated antigens (TAAs) or tumor-specific antigens (TSAs), strategies are required to improve delivery vehicles, optimize routes and timing of administration, and identify effective adjuvants to activate immune cells.^2^ To this end, biomaterial platforms, such as cryogels (a subclass of hydrogels), have been employed in preclinical models targeting melanoma, breast cancer, and other cancer subtypes.^20–30^ These scaffold-based vaccine platforms offer a promising solution through the protection of biomolecules from rapid clearance and degradation.^31–33^ The macroporous structure of cryogel scaffolds provides several benefits for the delivery of therapeutic cancer vaccines, including biocompatibility, mechanical strength, and injectability. The porous structure also facilitates nutrient diffusion and cell infiltration, providing an avenue to recruit and activate immune cells in situ.^25,34–36^ Cell-based therapies, TAA/TSA proteins or peptides, various adjuvants, and cytokines can be physically encapsulated or conjugated within the scaffold, along with granulocyte-macrophage colony-stimulating factor (GM-CSF), to recruit and stimulate antigen-presenting cells (APCs), such as DCs, at the injection site.^22,24,25,36^ Scaffold-based delivery of therapeutic cancer vaccines enhances the ability of the antitumor immune response to recognize and eliminate tumor cells by extending antigen release from the scaffold and allowing for effective antigen capture and activation of DCs to prime cellular (e.g., cytotoxic T cells) and humoral (e.g., antibody-dependent cellular cytotoxicity (ADCC)) responses.^36–39^

Loading efficiency and release kinetics of vaccine antigens and adjuvants have been a focus of improvement in scaffold-based delivery vehicles. Recombinant proteins typically exhibit rapid, diffusion-mediated burst release when physically encapsulated in hydrogels, including cryogel scaffolds.^40,41^ Extended vaccine exposure has been established as necessary to achieve the potency and breadth of immune responses required for adaptive immunity in targeting challenging diseases, such as cancer.^42–44^ Strategies to enhance protein antigen loading and release include implementing pre-absorbed protein cargo on Laponite nanoparticles to increase loading of protein and provide sustained release from an alginate hydrogel.^40^ Alternatively, tuning the viscoelastic properties of a poly(vinyl alcohol)-based hydrogel to promote sustained release of protein cargo has been recently explored.^45^ However, the ability of such strategies to enhance immunological outcomes has not been tested. Moreover, biorthogonal crosslinked hyaluronic acid (HA)-based cryogels supported sustained release of vaccine components resulting in increased intensity of the adaptive immune response over bolus injection but was only evaluated with highly immunogenic model antigen, ovalbumin (OVA) in a melanoma (B16-OVA) model.^23^ Sustained exposure of antigen was also recently shown through carbodiimide-mediated crosslinking of OVA peptides to create porous antigen-conjugated scaffolds which were proven to support sustained release of peptide through polymer hydrolysis and successfully activate antigen specific immune cells.^46^

Despite improvements in antigen delivery, these platforms remain limited by physiological immunosuppression and tolerance mechanisms. Subcutaneous scaffolds experience local hypoxia (0–3 % O_2_ in tissues) due to poor vascularization and high cellular activity during infiltration and innate immune response.^47–49^ In inflamed and cancerous tissues, hypoxia-driven signaling promotes the accumulation of extracellular adenosine, which potently inhibits effector functions of antitumor immune cells.^50–53^ Our previous work has shown that oxygenation of the TME (via respiratory hyperoxia and/or oxygenation agents) reduces intratumoral hypoxia and immunosuppressive extracellular adenosine while reprogramming the cytokine/chemokine/metabolome profile away from immunosuppression and toward immune-reactivity.^17,18,54,55^ Here, we apply the discovery that oxygenation potently reverses immunosuppression in the TME to our cryogel-based vaccine development. We previously engineered a HA-glycidyl methacrylate (HAGM) cryogel-based O_2_ delivery platform (O_2_-cryogel) that generates O_2_ and restores the function of DCs and T cells under hypoxic conditions.^56–58^ We have also shown the benefit of localized O_2_ availability and cryogel-based delivery in improving the efficacy of a protein subunit vaccine against COVID-19 (O_2_-Cryogel_VAX_).^59^ Local O_2_ generation within the scaffold can mitigate local hypoxia and oxygenate infiltrating cells, thereby functioning as a powerful co-adjuvant and reinforcing DC-mediated responses by preventing vaccine-associated hypoxia at the vaccination site.^56,59^ O_2_-Cryogel_VAX_ promoted a stronger humoral immune response, showing an increase in long-lasting anti-SARS-CoV-2 IgG antibodies compared to Cryogel_VAX_ and Bolus_VAX_. Additionally, O_2_ generation promoted the production of Th1-biased and pro-inflammatory cytokines.

Herein, we explored how enhanced antigen loading and release, in combination with local O_2_ generation, could enhance our Cryogel_VAX_ and O_2_-Cryogel_VAX_ platforms as effective delivery vehicles for a subunit protein cancer vaccine targeting prostate cancer. First, we explored the effectiveness of click chemistry (Click Cryogel_VAX_) via copper-free strain-promoted azide-alkyne cycloaddition (SPAAC) to promote the delivery of a TAA protein, specifically prostatic acid phosphatase (PAP). SPAAC-mediated conjugation has been shown to be a safe biomolecule-conjugation approach in HAGM-based cryogel scaffolds.^60^ Click Cryogel_VAX_ was evaluated against Cryogel_VAX_ to determine how extended exposure of TAA to infiltrating APCs might enhance B-and T-cell responses post-immunization. Subsequently, local oxygenation was further evaluated as a potent co-adjuvant (O_2_-Cryogel_VAX,_ Click O_2_-Cryogel_VAX_). Finally, we evaluated the efficacy of Click O_2_-Cryogel_VAX_ to elicit a strong, balanced immune response against the loaded TAA, thereby supporting an effective antitumor response in a TRAMP-C2 prostate cancer model.

## Results

### Fabrication of prostate cancer-associated antigen cryogel-based vaccines

To engineer a prostate cancer vaccine that elicits immunologic memory against murine prostate cancer cells (TRAMP-C2), the overexpressed protein prostatic acid phosphatase (PAP) was used in addition to the immunomodulatory factors; granulocyte-macrophage colony-stimulating factor (GM-CSF) and polyinosinic:polycytidylic acid (Poly(I:C)). Previous cryogel-based vaccine formulations relied on physical entrapment of antigens and adjuvants during cryogelation to deliver the vaccine. Here, we found that PAP exhibits poor loading through physical encapsulation, therefore we evaluated if chemical conjugation of PAP through click bioconjugation would influence vaccine efficacy. First, the polymer, HAGM, was modified with Azido-PEG3-amine through carbodiimide (EDC/NHS) coupling, and the resulting polymer HAGM-PEG3-Azido was purified. Conjugation of the Azido-PEG3 group to the HAGM backbone was confirmed with ^1^H NMR and ATR-FTIR analysis **(Figs. S1, S2)**. Additionally, PAP was reacted with DBCO-PEG4-NHS ester to create DBCO-PEG4-PAP and purified. HAGM or HAGM-PEG3-Azido were combined with PAP or DBCO-PEG4-PAP, respectively, and GM-CSF and Poly(I:C) in solution overnight at 4 °C before cryopolymerization at -20 °C for 18 h and then thawed to form cryogel-based vaccines, Cryogel_VAX_ (HAGM) or Click Cryogel_VAX_ (HAGM-PEG3-Azido) **(Fig. 1A)**. To form O_2_-generating cryogel-based vaccines, ball-milled calcium peroxide (CaO_2_) particles (1 % w/v) and acrylate-PEG-catalase (CAT) (1 % w/v) were incorporated into the polymer solution before freezing **(Fig. 1A)**. CaO_2_ hydrolyzes with water (H_2_O) to form hydrogen peroxide (H_2_O_2_) and O_2_. CAT is added to support the decomposition of H_2_O_2_ into H_2_O and O_2_.

**Figure 1.**
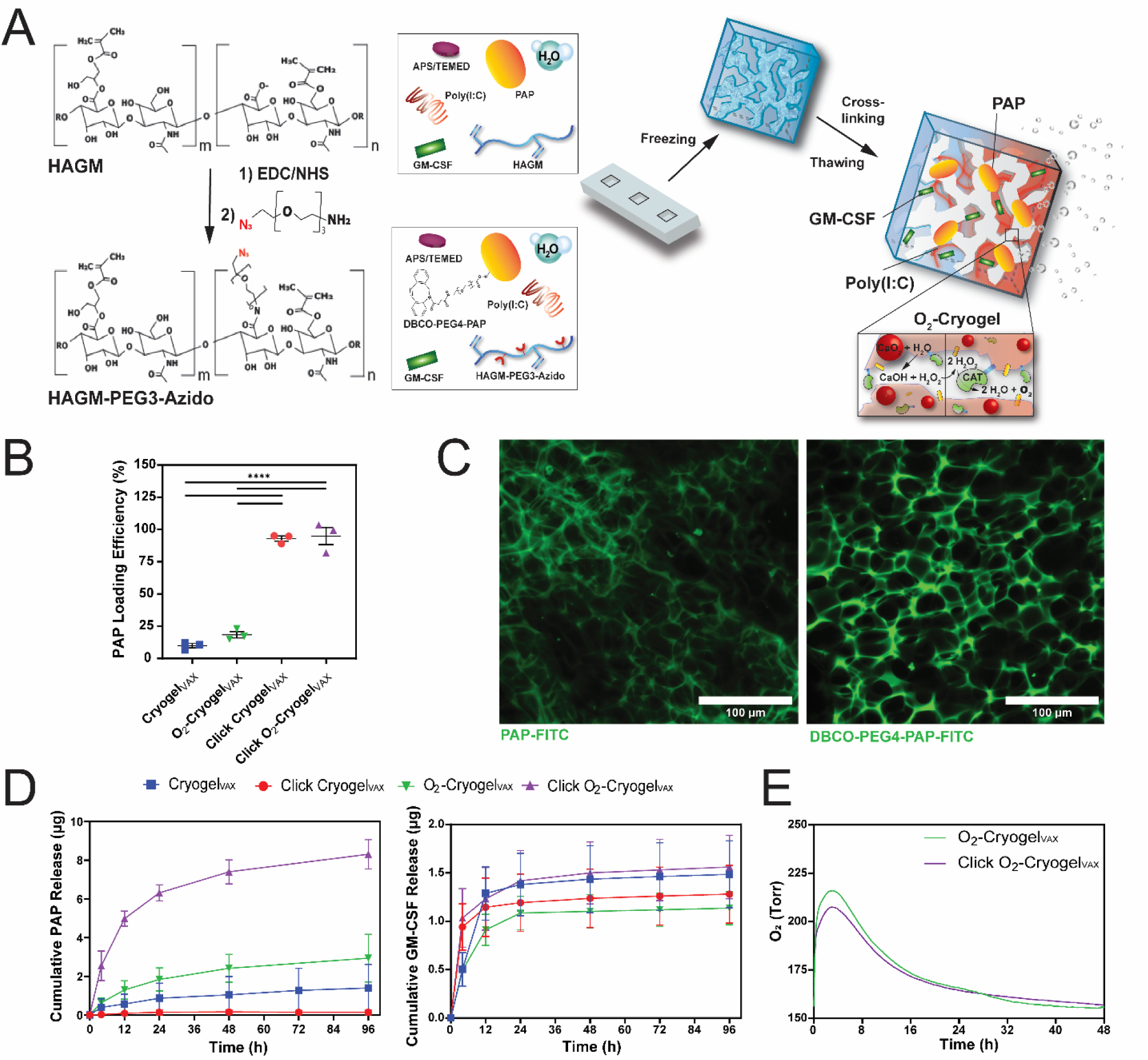
Characterization of PAP-containing cryogel-based vaccines. **(A)** Schematic depicting the fabrication of Cryogel_VAX_ and O_2_-Cryogel_VAX_ using HAGM or HAGM-PEG3-Azido, prostate cancer antigenic proteins, and O_2_ generation. In this process, the polymer is dissolved with biomolecules and calcium peroxide (CaO_2_) before being mixed with the initiator system (APS/TEMED), followed by incubation at −20 °C for 18 h to allow cryopolymerization. **(B)** Loading efficiency (%) of PAP inside cryogel-based vaccines. Values are presented as mean ± SEM of three independent experiments. **(C)** Representative confocal microscopy images of FITC-conjugated PAP and DBCO-PEG4-PAP inside Cryogel_VAX_ and Click Cryogel_VAX_. **(D)** In vitro release curves for PAP and GM-CSF from cryogel vaccines over time. Values are presented as mean ± SD (n = 3–4). **(E)** O_2_ release from O_2_-Cryogel_VAX_ and Click O_2_-Cryogel_VAX_ over time. Values are presented as mean (n = 3). Statistical analysis was performed using one-way ANOVA and a Bonferroni post hoc test; ****p < 0.0001.

Encapsulation of PAP through physical entrapment resulted in poor loading efficiency, 10 ± 1.7 % in Cryogel_VAX_ and 18 ± 2.4 % in O_2_-Cryogel_VAX_. As expected, click conjugation ensured nearly complete loading of the DBCO-modified protein with 93 ± 2.0 % and 95 ± 6.6 % loading efficiency in Click Cryogel_VAX_ and Click O_2_-Cryogel_VAX_, respectively **(Fig. 1B).** Confocal microscopy of FITC-labeled protein confirmed protein within the polymer walls of macroporous cryogel scaffolds **(Fig. 1C)**. PAP was released over 96 h, with full recovery of the encapsulated protein from Cryogel_VAX_ and O_2_-Cryogel_VAX_. As expected, chemical conjugation prevented release of PAP from Click Cryogel_VAX_; however, almost all of the encapsulated PAP (∼90 %) was released from Click O_2_-Cryogel_VAX_ within 96 h **(Fig. 1D).** This finding is attributed to degradation-induced release in O_2_-Cryogel_VAX_. This is consistent with our previously reported findings that the introduction of O_2_-generation into the cryogel supports oxidation-mediated degradation, which suggests that PEG and HAGM degradation allows for the release of PAP over time.^57,61–63^ Similar to previously reported findings, GM-CSF was physically entrapped within Cryogel_VAX_, 99 ± 23 %, and was released within 12–24 h. No significant differences were found with the addition of click moieties and O_2_ generation **(Fig. 1D).**

O_2_-generating cryogel vaccines released O_2_ for 48 h, and the O_2_ level peaked around 3 h at 200–225 Torr. Both O_2_-Cryogel_VAX_ and Click O_2_-Cryogel_VAX_ exhibited very similar release profiles for O_2_ **(Fig. 1E).** To confirm no toxicity risk from H_2_O_2_ production and decomposition by CAT, O_2_-Cryogel_VAX_ and Click O_2_-Cryogel_VAX_ were incubated in PBS with and without covalently included CAT in the scaffold, and the released H_2_O_2_ in the supernatant was measured. As expected, the inclusion of CAT (1 % w/v) was able to deplete the H_2_O_2_, maintaining safe levels below 10 µM, similar to Cryogel alone, compared to CAT-free O_2_-cryogels which produced ∼30 µM H_2_O_2_ **(Fig. S3).**

### Immunization with Click O_2_-Cryogel_VAX_ improves APC response in draining lymph nodes (LNs)

We hypothesized that APCs would traffic to the injection site, become activated and migrate to the draining LNs to prime effective B- and T-cell responses against the delivered antigen **(Fig. 2A).** To test if the Cryogel_VAX_ formulations were effective at promoting immune-cell responses in the draining LNs, 6–8-week-old male C57BL/6 mice were immunized and boosted at day 0 (D0) and 14 (D14) with subcutaneous injection on both flanks. Mice were euthanized on day 28 (28D) after initial immunization, and the inguinal LNs were dissected for evaluation of APC presence through flow cytometry **(Fig. S4A)**. Vaccination with Click O_2_-Cryogel_VAX_ created the strongest inflammatory response with the highest number of CD45^+^ leukocytes **(Fig. S4B)**. Click O_2_-Cryogel_VAX_ significantly increased the total count of CD11c^+^ MHC-II^+^ conventional dendritic cells (cDCs) compared to Bolus_VAX_ injection (p = 0.0064) and Cryogel_VAX_ (p = 0.0156) **(Fig. 2B)**. Additionally, Click O_2_-Cryogel_VAX_ significantly increased the total count of F4/80^+^ macrophages compared to Bolus_VAX_ (p = 0.0005), O_2_-Cryogel (p = 0.0035), and Cryogel_VAX_ (p = 0.0033) **(Fig. 2C)**.

**Figure 2.**
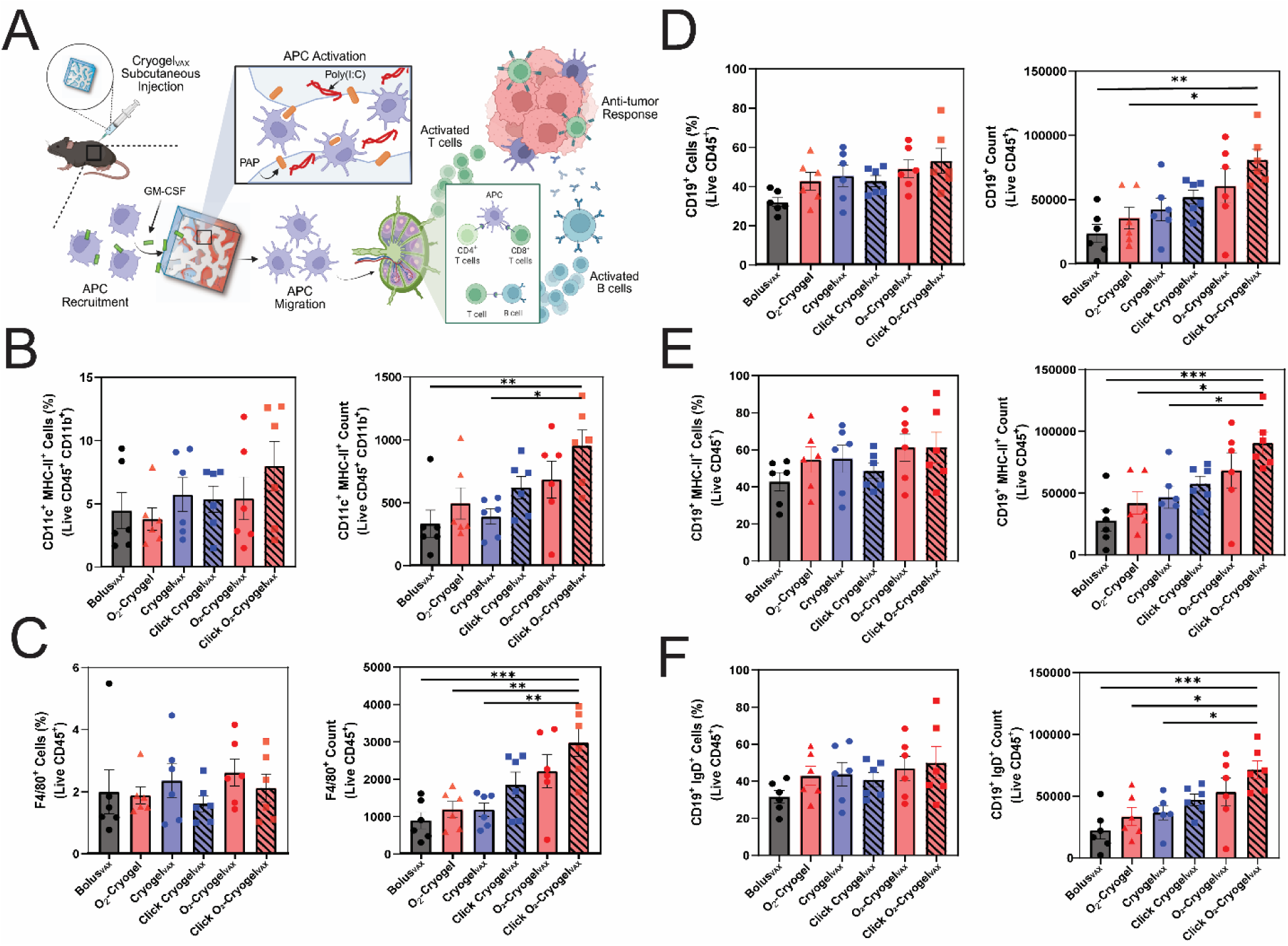
Vaccination with Click O_2_-Cryogel_VAX_ promotes APC expansion in the LNs. **(A)** Schematic illustrating cryogel-based vaccine supported immunity through recruitment of APCs that will migrate to the LNs and prime antitumor T and B cell responses. Created with BioRender.com **(B)** Percentage and count of conventional DCs (CD11c^+^, MHC-II^+^) in inguinal LNs. **(C)** Percentage and count of macrophages (F4/80^+^) in inguinal LNs. **(D)** Percentage and count of B cells (CD19^+^) in inguinal LNs. **(E)** Percentage and count of antigen-presenting B cells (CD19^+^ MHC-II^+^) in inguinal LNs. **(F)** Percentage and count of mature B cells (CD19^+^ IgD^+^) in inguinal LNs. Two draining LNs were analyzed per animal. Values are indicative of the average from each animal and are presented as mean ± SEM (n = 6). Statistical analysis was performed using one-way ANOVA and Bonferroni post hoc test: ^∗^p < 0.05, ^∗∗^p < 0.01, and ^∗∗∗^p < 0.001.

Next, the vaccine’s ability to enhance B-cell activity in the LNs was evaluated. Click O_2_-Cryogel_VAX_ significantly increased the number of CD19^+^ B cells over Bolus_VAX_ (p = 0.0012) and O_2_-Cryogel (p = 0.0144) **(Fig. 2D)**. An increase in the number of antigen-presenting MHC-II^+^ B cells was observed with Click O_2_-Cryogel_VAX,_ showing the best response over Bolus_VAX_ (p = 0.0008), O_2_-Cryogel (p = 0.0133), and Cryogel_VAX_ (p = 0.0319) **(Fig. 2E)**. Additionally, Click O_2_-Cryogel_VAX_ increased total counts of mature IgD^+^ B cells compared to Bolus_VAX_ (p = 0.0009), O_2_-Cryogel (p = 0.0142), and Cryogel_VAX_ (p = 0.0297) **(Fig. 2F).** These results suggest that Click O_2_-Cryogel_VAX_ exhibits a higher APC prevalence in the draining LNs than previous cryogel vaccine formulations and Bolus vaccination.

### O_2_ generation increases Th1 antibody response and improves TAA-specific antibody production

To determine whether the increased B-cell presence after vaccination was effective in eliciting a robust antibody response, blood was collected from the submandibular vein of the mice 2 weeks after the prime and boost on D14 and D28, and the serum was isolated **(Fig. 3A)**. Antibody titers were found through serial dilution of mouse sera and evaluated using Immunoglobulin isotyping (LegendPlex) and ELISA. Antibody potency was plotted and fitted to a 4-parameter logistic nonlinear regression curve, and the EC50 (half-maximal effective concentration) was calculated as the antibody titer for each treatment group **(Fig. S5)**. As expected, inclusion of O_2_-generation improved the overall humoral immune response. Click O_2_-Cryogel_VAX_ significantly increased PAP-specific immunoglobulin production by D28 over Bolus_VAX_ (p = 0.0078), O_2_-Cryogel alone (p = 0.0093), Cryogel_VAX_ (p = 0.0083), and Click Cryogel_VAX_ (p = 0.009), additionally, while not statistically significant, O_2_-Cryogel_VAX_ also showed a similar increase in PAP-specific immunoglobulin at D28 **(Fig. 3B)**. Immunization with Click O_2_-Cryogel_VAX_ also significantly increased the levels of primary antibody IgM at D28 compared to Bolus_VAX_ (p = 0.0285), Cryogel_VAX_ (p = 0.0155), and Click Cryogel_VAX_ (p = 0.0063) **(Fig. 3C)**.

**Figure 3.**
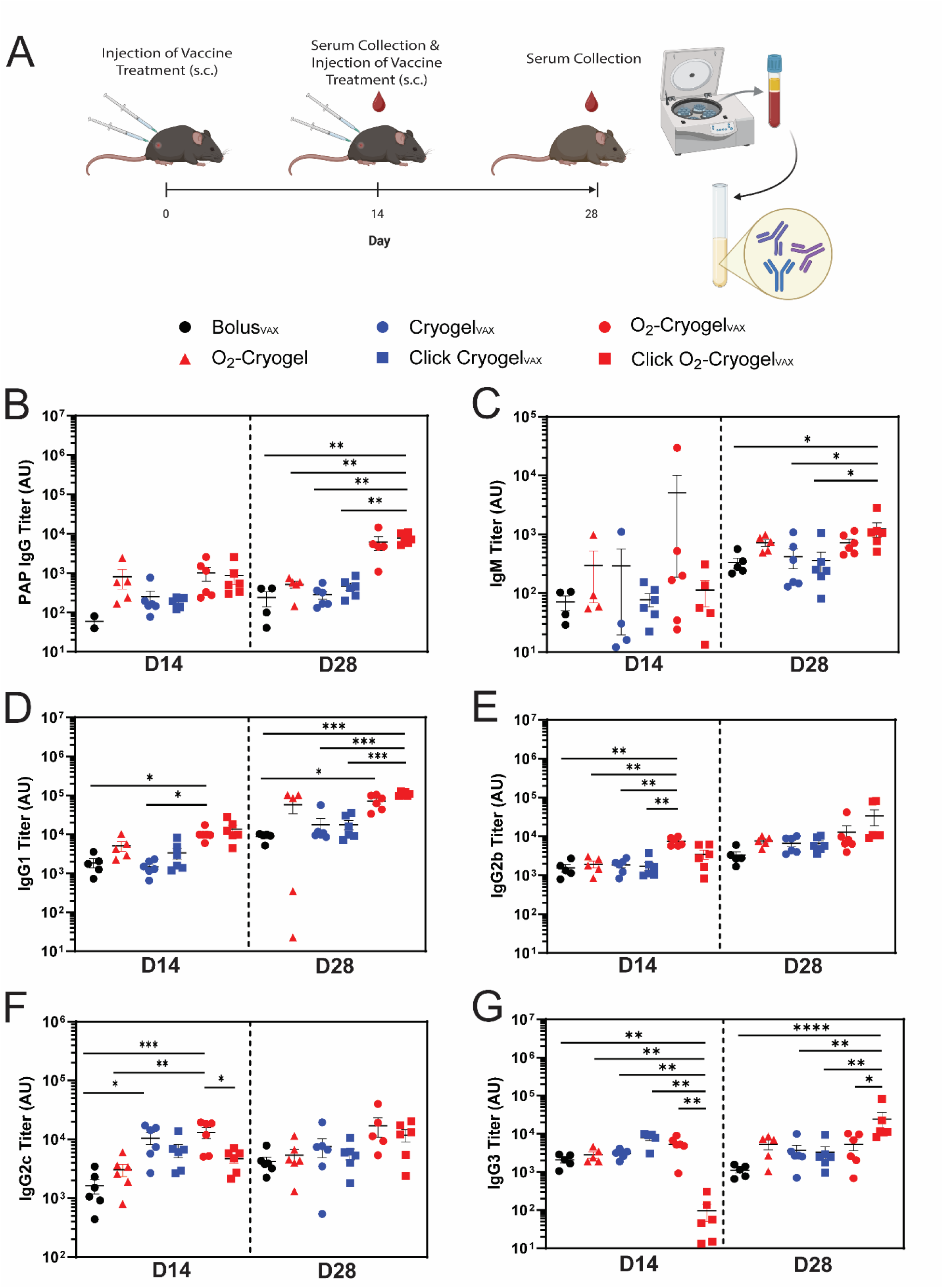
O_2_-generating cryogel-based vaccines promote production of Th1-associated antibodies. **(A)** C57BL/6 mice were immunized subcutaneously on day 0 (D0) and day 14 (D14). Mouse sera was collected 14 d after each immunization and analyzed for anti-PAP antibodies and IgG subtypes. Schematic created with BioRender.com **(B)** Anti-PAP IgG, **(C)** IgM, **(D)** IgG1, **(E)** IgG2b, **(F)** IgG2c, **(G)** IgG3 EC50 titers on D14 and day 28 (D28) after prime and boost with Bolus_VAX_, O_2_-Cryogel alone or Cryogel_VAX_ formulations. Values are presented as mean ± SEM (n = 5–6). Statistical analysis was performed using one-way ANOVA and Bonferroni post hoc test: *p < 0.05, **p < 0.01, ***p < 0.001, and ****p < 0.0001.

To better understand the balance between Th1/Th2 immune responses after vaccination, the overall production of several IgG subclasses was evaluated. Mice that received O_2_-Cryogel_VAX_ or Click O_2_-Cryogel_VAX_ had elevated IgG1 titers on D14 and D28 **(Fig. 3D)**. While there were no significant differences on D28, O_2_-Cryogel_VAX_ significantly increased the levels of IgG2b and IgG2c on D14, and a similar tendency is observed on D28 **(Fig. 3E–F)**. However, vaccination with Cryogel_VAX_ and Click Cryogel_VAX_ displayed similar increase in IgG2c as with oxygenation **(Fig. 3F)**. Interestingly, we found that Click O_2_-Cryogel_VAX_ had significantly lower levels of IgG3 on D14 that was overcome by D28 with significantly higher levels compared to the other treatments **(Fig. 3G)**. These findings suggest that cryogel delivery of the vaccine components supports Th1 cells, indicated through increased levels of IgG2c, while O_2_ generation increased levels of IgG1 and IgG3, suggesting it could help promote antibody-dependent cellular cytotoxicity (ADCC), as IgG1 and IgG3 target the Fc-dependent (FcγRIIIa) effector functions of NK cells in targeting cancer cells.^64^

### Improved loading and delivery of TAA support memory cell generation and cytotoxic responses

To evaluate the cellular immune response after vaccination, the spleens were dissected after 28 d and memory T cell populations were characterized with flow cytometry **(Fig. S6)**. Strikingly, Cryogel_VAX_ and Click Cryogel_VAX_ resulted in a significant shift in the proportion of central memory (CD44^+^ CD62L^+^) T cells over effector memory (CD44^+^ CD62L^-^) in both CD4^+^ and CD8^+^ T cells, suggesting that cryogel vaccines supported the creation of long-lasting immunity compared to Bolus_VAX_ **(Fig. 4A–F)**. Cryogel_VAX_ and Click Cryogel_VAX_ increased CD4^+^ central memory T cells over Bolus_VAX_ (p = 0.0081, 0.0016) and Click O_2_-Cryogel_VAX_ maintained a significant increase compared to Bolus_VAX_ (p = 0.0185) **(Fig. 4B)**. Similarly, in the CD8^+^ central memory T cells, Cryogel_VAX_ and Click Cryogel_VAX_ increased over Bolus_VAX_ (p = 0.0099, 0.0018), and Click O_2_-Cryogel_VAX_ maintained a significant increase compared to Bolus_VAX_ (p = 0.0044) **(Fig. 4E)**. Interestingly, the increase in central memory population was maintained, or potentially slightly attenuated, by the inclusion of O_2_ generation, suggesting that cryogel-supported vaccine delivery plays a central role in the generation of memory T cells. Additionally, Cryogel_VAX_, Click Cryogel_VAX_, and Click O_2_-Cryogel_VAX_ significantly increased the population of Naïve T cells, indicating an altered proportion of T-cell population after vaccination **(Fig. 4C**, **4F)**.

**Figure 4.**
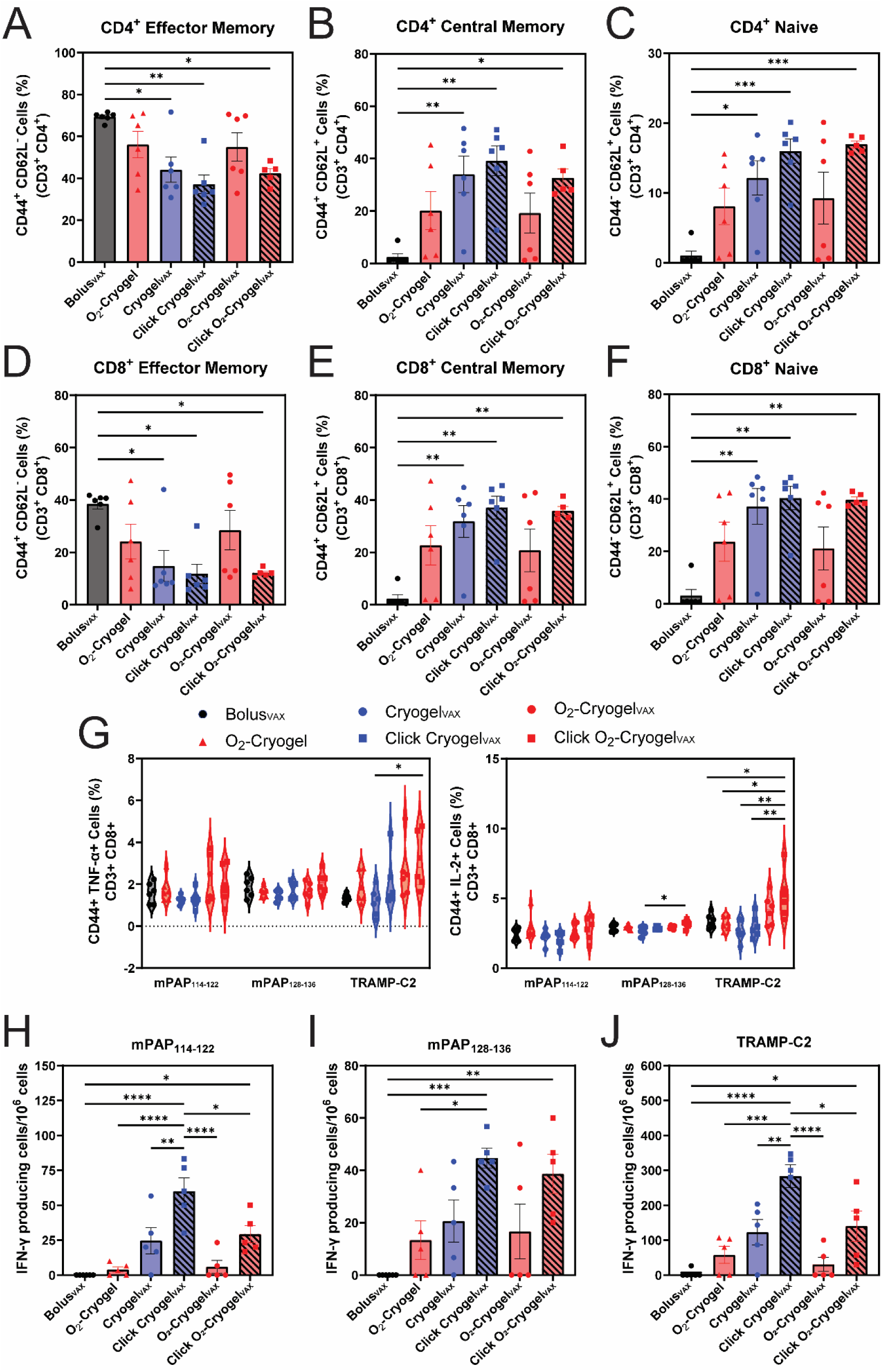
Vaccination with Click Cryogel_VAX_ and Click O_2_-Cryogel_VAX_ promote the generation of robust cellular memory to tumor-associated antigens. **(A–F)** Flow cytometric analysis of memory T cell populations in spleen after receiving vaccine regimen including percentages of CD4^+^ **(A)** effector memory (CD44^+^, CD62L^-^), **(B)** central memory (CD44^+^, CD62L^+^), and **(C)** naïve (CD44^-^, CD62L^+^), and percentages of CD8^+^ **(D)** effector memory (CD44^+^, CD62L^-^), **(E)** central memory (CD44^+^, CD62L^+^), and **(F)** naïve (CD44^-^, CD62L^+^). **(G)** Percentages of TNF-α-producing (CD44+, TNF-α+) and IL-2-producing (CD44^+^, IL-2^+^) activated CD8^+^ T cells in response to splenocyte restimulation with TAAs. **(H–J)** TAA specific IFN-γ producing splenocytes, as a proxy for T cells after culture with peptides and tumor cells. Values are presented as mean ± SEM (n = 5–6). Statistical analysis was performed using one-way ANOVA and Bonferroni post hoc test: *p < 0.05, **p < 0.01, ***p < 0.001, and ****p < 0.0001.

To evaluate TAA-specific memory, the collected splenocytes were co-cultured with TAA-peptides (mPAP_114-122_, mPAP_128-136_), recombinant PAP, TRAMP-C2 cells, PMA/ionomycin (positive-control), or culture media alone (negative-control) for 6 h, and the cellular immune response was evaluated by measuring the generation of Th1 cytokines: TNF-α, IL-2, and IFN-γ. Vaccination with Click O_2_-Cryogel_VAX_ significantly increased the percentage of CD44^+^ TNF- α^+^ CD8^+^ T cells in response to the tumor cells **(Fig. 4G, Fig. S7)** and the percentage of CD44^+^ IL-2^+^ CD8^+^ T cells in response to the tumor cells and TAA peptide mPAP_128-136_ **(Fig. 4G, Fig. S7)**. Splenocytes collected from mice that received Click Cryogel_VAX_ and Click O_2_-Cryogel_VAX_ additionally produced more IFN-γ when exposed to TAA peptides mPAP_114-122_ and mPAP_128-136_ as well as the tumor cells **(Fig. 4H–J)**. These results suggest reinforced anti-cancer outcomes with click-loaded protein by increased production of Th1 cytokines, which support the differentiation and activation of cytotoxic T cells, macrophages, and natural killer cells.

### O_2_-Cryogel_VAX_ and Click O_2_-Cryogel_VAX_ significantly reduce tumor growth in a therapeutic TRAMP-C2 model

Based on our results above, demonstrating that click-mediated loading improved cellular memory after vaccination and that O_2_-generation could boost ADCC and Th1 responses, we evaluated the anti-cancer therapeutic efficacy of Click O_2_-Cryogel_VAX_. TRAMP-C2 cells were derived from a mouse prostate adenocarcinoma and is considered an immunologically “cold” tumor model.^65,66^ TRAMP-C2 tumors were subcutaneously developed in 8–10-week-old male C57BL/6 mice. When the mean tumor size exceeded 50 mm^3^, mice received a prime (0 d) and boost (14 d) immunization of Bolus_VAX_, O_2_-Cryogel, O_2_-Cryogel_VAX_, or Click O_2_-Cryogel_VAX_ in both flanks, and tumor size and mouse survival was monitored over time **(Fig. 5A)**. As compared to our control, Bolus_VAX_, the use of O_2_-cryogels, as described before, slowed tumor progression and slightly increased survival by 2 d. Moreover, when combined with TAA, O_2_-Cryogel_VAX_ significantly hindered tumor growth compared to Bolus_VAX_ (p = 0.0005) and O_2_-Cryogel alone (p = 0.0003) **(Fig. 5B–C)**. Notably, improved loading of the TAA in Click O_2_-Cryogel_VAX_ led to an enhanced antitumor response with a significant decrease in tumor growth rate. Treatment with Click O_2_-Cryogel_VAX_ significantly slowed tumor growth compared to Bolus_VAX_ (p = 0.0005) and O_2_-Cryogel alone (p < 0.0001) **(Fig. 5B–C)**.

**Figure 5.**
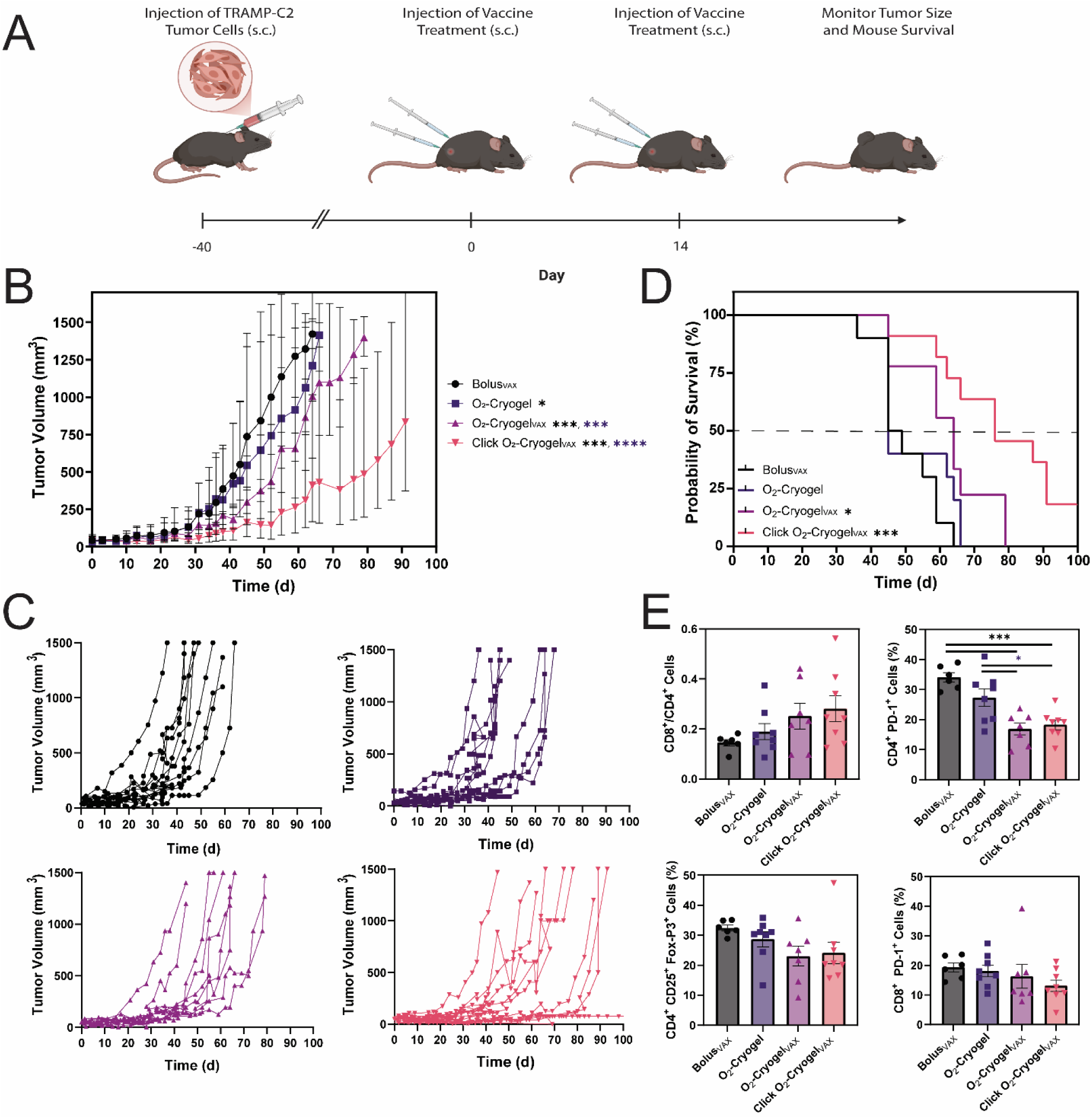
Vaccination with Click O_2_-Cryogel_VAX_ significantly delays TRAMP-C2 tumor growth. **(A)** C57BL/6 mice were subcutaneously injected with TRAMP-C2 tumor cells. After 40 d, mice were immunized and boosted 2 weeks apart, and tumors were measured with calipers until tumor burden exceeded euthanasia criteria. Schematic created with BioRender.com **(B)** Average tumor volume over time post-treatment and **(C)** individual subject tumor volume over time post-treatment with Bolus_VAX_, O_2_-Cryogel, O_2_-Cryogel_VAX_, and Click O_2_-Cryogel_VAX_. Data are presented as mean ± SEM (n = 10–11). Statistical analysis was performed using Mixed effects analysis and Bonferroni multiple comparisons test: *p < 0.05, ***p < 0.001, and ****p < 0.0001. **(D)** Overall Kaplan–Meier survival curves of treated mice (n = 10–11). Statistical analysis was performed using log-rank (Mantel-Cox) test and Bonferroni multiple comparisons test: *p < 0.05, and ***p < 0.001. **(E)** Flow cytometric characterization of tumor-infiltrated T cells. Data are presented as mean ± SEM (n = 6–8). Statistical analysis was performed using one-way ANOVA and Bonferroni post hoc test: *p < 0.05, and ***p < 0.001. Black colored * indicates comparison to Bolus_VAX_, and purple colored * indicates comparison to O_2_-Cryogel.

Strikingly, treatment with O_2_-Cryogel_VAX_ and Click O_2_-Cryogel_VAX_ increased the median survival to 64 and 76 d, respectively, compared to 47 d with Bolus_VAX_ (p = 0.026, p = 0.0004) **(Fig. 5D)**. The median lifespan of mice receiving Click O_2_-Cryogel_VAX_ was extended by 62 %. In addition, Click O_2_-Cryogel_VAX_ virtually ceased tumor growth immediately after treatment for 2 mice until study endpoint. When characterizing the tumor-infiltrating T cells **(Fig. S8)**, O_2_-Cryogel_VAX_ and Click O_2_-Cryogel_VAX_ incrementally increased the presence of CD8^+^ tumor-infiltrating T cells, resulting in an increased ratio of CD8^+^/CD4^+^ T cells **(Fig. 5E)**. While not statistically significant both O_2_-Cryogel_VAX_ (23.1 ± 3.3 %) and Click O_2_-Cryogel_VAX_ (24.1 ± 3.6 %) similarly decreased the percentage of CD25^+^ FoxP3^+^ regulatory T cells compared to less effective treatments Bolus_VAX_ (32.4 ± 0.95 %) and O_2_-Cryogel (28.7 ± 2.5 %) **(Fig. 5E).** Furthermore, O_2_-Cryogel_VAX_ (16.9 ± 2.0 %) and Click O_2_-Cryogel_VAX_ (18.3 ± 1.6 %) significantly decreased the percentage of PD-1^+^ CD4^+^ cells over Bolus (34.1 ± 1.6 %) and O_2_-Cryogel (27.3 ± 2.9 %) (p < 0.001, p < 0.05) **(Fig. 5E)**. A similar trend was also obtained in PD-1^+^ CD8^+^ cells. Taken together, these results suggest that vaccination with O_2_-Cryogel_VAX_ and Click O_2_-Cryogel_VAX_ promote antitumor T-cell responses resulting in significant prostate tumor regression.

## Discussion

Recent advancements in vaccine development provide promising opportunities to enhance the design and delivery of effective therapeutic cancer vaccines.^67,68^ However, creating an effective adaptive immune response that can overcome barriers to cancer therapy, such as the immunosuppressive TME, tolerance to self-antigens, and tumor heterogeneity, has proven challenging.^67,68^ Harnessing biomaterial scaffolds, such as cryogels, as a delivery platform may be key to enhancing vaccine efficacy.^22,24,27,36,38^ By delaying the release of vaccine components, scaffold-based delivery enables improved adaptive memory responses after immunization.^29,37,40,42^ Additionally, we demonstrated that O_2_ delivery can be utilized as a co-adjuvant to boost antitumor responses to overcome physiological immune suppression and have incorporated this effective strategy into our vaccine design.^56,59,69,70^ The enhanced immune responses observed may be explained by restoration of DC function under hypoxic conditions, as previously shown.^56,58^ O₂-cryogels prevent tolerogenic DC phenotypes and restore antigen presentation and T-cell priming. We leveraged click-mediated covalent bioconjugation to deliver the subunit antigenic protein within a cryogel-based vaccine platform to enhance adaptive memory and utilized O_2_ as a powerful co-adjuvant to boost the Th1 proinflammatory response. Together these two unique implementations to our vaccine platform promoted a strong antitumor response.

To formulate a Cryogel_VAX_ platform for murine prostate cancer, we incorporated TAA protein PAP, along with recombinant murine GM-CSF to recruit APCs to the cryogel and the adjuvant Poly(I:C) to activate APCs against the self-antigen. The biopolymer HAGM was selected for its demonstrated performance in the formulation of biocompatible, injectable cryogels and was previously utilized in Cryogel_VAX_ and O_2_-Cryogel_VAX_.^59,60,71^ Additionally, HA-based polymers are biodegradable compared to alginate or synthetic polymers.^61,72,73^ However, the loading efficiency of PAP in the HAGM-based cryogel was low; therefore, we investigated how covalent bioconjugation of PAP via click chemistry would influence the immune response after immunization. By integrating an azido-group into the HAGM backbone and adding a DBCO-group to the protein antigen, we were able to encapsulate 100 % of the dose within the cryogel-scaffold. This is paramount to increasing the clinical translatability of the delivery vehicle, as it prevents the undesired waste of biomolecules during formulation. SPAAC chemistry was chosen as it has already been shown as a biocompatible and effective strategy to conjugate biomaterials using click chemistry into HAGM cryogel.^60^ However, we altered the strategy to conjugate the protein prior to cryopolymerization to ensure compatibility with O_2_-release and incorporated PEG linkers to encourage specific receptor-ligand interaction.^74^ Future studies should be conducted to investigate the influence of linker length on vaccine efficacy. Additionally, PEG has been suggested to be immunogenic, therefore it might influence the activation of infiltrating APCs.^75^ Furthermore, O_2_ generation likely induces oxidative degradation of the cryogel matrix, enabling controlled release of covalently bound antigens, as supported by prior work.^57^ This dual mechanism combines sustained antigen retention with environment-triggered release.

Surprisingly, we found that improved TAA delivery in Click Cryogel_VAX_ did not increase the production of anti-PAP antibodies nor change the presentation or levels of antibody subtypes compared to Cryogel_VAX_ nor our control Bolus_VAX_. However, O_2_ delivery was consistent between O_2_-Cryogel_VAX_ and Click O_2_-Cryogel_VAX_, in increasing IgG1 titers as previously shown with O_2_-Cryogel_VAX_ for COVID-19.^59^ Additionally, we showed that cryogel delivery skewed stronger IgG2c titers, but our findings contrast with previous studies showing that HA-based cryogel delivery improved IgG1 and IgG2b titers; although these studies used strongly immunogenic proteins and CpG-ODN as an adjuvant.^23,59^ It would be necessary to further investigate the adjuvant effect in skewing the immune response. However, Click O_2_-Cryogel_VAX_ showed a potential synergistic effect, resulting in the highest levels of PAP-specific binding antibodies and overall IgM, IgG1, IgG2b, and IgG3 antibodies, suggesting that both antigen dose and O_2_ were key in skewing a Th1 antibody response.

Immunization with Cryogel_VAX_ and Click Cryogel_VAX_ skewed the generation of central memory T cells over effector memory T cells in both CD4^+^ and CD8^+^ T cells in the spleen. This finding suggests that scaffold-based delivery plays an important role in generating central memory relative to Bolus_VAX_. Unexpectedly, while Click Cryogel_VAX_ slightly increased the mean percentages of central memory T cells over Cryogel_VAX_ it was not a significant increase, suggesting that protein dose may not have influenced the general memory profile. However, our results support previous studies that reported sustained release of antigen with rapid release of adjuvants can drive Th1 responses.^37^ When splenocytes were exposed to TAA-derived peptide epitopes and TRAMP-C2 cells, Click O_2_-Cryogel_VAX_ promoted the expression of Th1-biased proinflammatory cytokines, TNF-α and IL-2. Interestingly, Click Cryogel_VAX_ elicited the strongest IFN-γ response but was hindered by O_2_ production. This finding warrants further investigation into whether the decreased response is attributable to O_2_ generation or to faster antigen release from Click O_2_-Cryogel_VAX_ compared with Click Cryogel_VAX_.

Optimizing Cryogel_VAX_ with click-mediated covalent conjugation to deliver the entire protein dose ensured enough exposure to the TAA, generated a TAA-specific response, and supported the development of central memory. Additionally, O_2_ release performed as a co-adjuvant to create a stronger Th1-biased response. The presence of central memory T cells is critical to the efficacy of anti-cancer immunotherapies, and a robust Th1 response is essential for mounting an effective, durable cellular attack against cancer.^76–78^ An increased presence of memory T cells has also been shown to improve therapeutic chimeric antigen receptor (CAR)-T cell therapy outcomes. CAR-T cell products enriched for naïve and early memory subsets correlate with superior CAR-T cell expansion, persistence, and efficacy.^79,80^ Using multiplex gene editing approaches, we recently demonstrated that CAR-T cells can be engineered to be resistant to hypoxia-driven biochemical and immunological negative regulators, thereby enabling an allogeneic CAR-T therapy that eliminates immunosuppressive solid tumors in humanized xenograft mouse model.^81^ Findings of this study suggest that treatment with cryogel-based vaccines could be an effective strategy to prime naïve and central memory anti-cancer T cells to synergize with CAR-T therapies, or be used to generate a source of memory or Th1 T cells to be used for CAR T-cell generation.

Finally, treatment with Click O_2_-Cryogel_VAX_ elicited a potent anticancer response in the immunologically cold TRAMP-C2 tumor model and extended the median lifespan of mice receiving Click O_2_-Cryogel_VAX_ by 62 %. These findings support Click O_2_-Cryogel_VAX_ potential as a more effective delivery vehicle for therapeutic cancer vaccines and could increase the immunogenicity of delivered neoantigens for personalized vaccines.^82^ Click-mediated conjugation enables platform modularity for various vaccine applications, as any antigenic protein or peptide can be readily substituted. Further research is needed to assess the duration of the vaccine-induced immune responses, and the optimal dosing required to maintain tumor-free survival. Future direction of this work will investigate the use of Click O_2_-Cryogel_VAX_ in combination with other immunotherapies, such as ICIs and ACT.

## Materials and Methods

### Materials

HA sodium salt (∼1.5 MDa), Dulbecco’s phosphate buffered saline (PBS), sodium bicarbonate (NaHCO_3_), MES hydrate (C_6_H_13_NO_4_S · xH_2_O), glycidyl methacrylate (GM), N, N-dimethylformamide (DMF), triethylamine (TEA), tetramethylethylenediamine (TEMED), ammonium persulfate (APS), N-(3-dimethylaminopropyl)-N′-ethylcarbodiimide (EDC), N-hydroxysuccinimide (NHS), fetal bovine serum, Catalase from Aspergillus niger, and Insulin solution from bovine pancreas were acquired from MilliporeSigma (St. Louis, MO, USA). Deuterium oxide (D_2_O), penicillin/streptomycin, (+)-Dehydroisoandrosterone, and eBioscience^TM^ Fixable Viability Dye eFluor^TM^ 506 were purchased from Thermo Fisher Scientific (Waltham, MA, USA). The TRAMP-C2 cells were purchased from American Type Culture Collection (Manassas, VA, USA). Recombinant murine granulocyte-macrophage colony-stimulating factor (GM-CSF), Dulbecco’s Modified Eagle Medium (DMEM), and Roswell Park Memorial Institute 1640 medium (RPMI) were procured from Gibco (Waltham, MA, USA). Poly(I:C) (HMW) VacciGrade™ was purchased from InvivoGen (San Diego, CA, USA). Acrylate-polyethylene glycol (PEG)-succinimidyl (AP-NHS, 3.4 kDa) was purchased from Laysan Bio (Arab, AL, USA). Recombinant murine PAP was purchased from Abcam (Cambridge, UK). Azido-PEG3-amine and DBCO-PEG4-NHS Ester were from Vector Labs (Newark, CA, USA).

### Methods

#### Synthesis of HAGM and HAGM-PEG3-Azido

First, HAGM was synthesized as previously reported.^83,84^ Briefly, HA (Sigma) reacted with GM in a co-solvent mixture of PBS and DMF for 10 d, and then subsequently precipitated in acetone and vacuum dried. To functionalize with an azido group, HAGM (1 % w/v) was dissolved in MES buffer (100 mM, pH 6.5). EDC and NHS were sequentially added to the polymer solution at 5 and 6 equivalents, respectively, and allowed to react for 15 minutes. Azido-PEG3-amine (Vector Labs) was dissolved in solution at a 1:10 molar ratio relative to carboxyl groups, and the reaction was conducted for 24 h protected from light. The polymer solution was transferred to dialysis tubing (10 kDa MWCO) and dialyzed against deionized H_2_O for 24 h, with the H_2_O changed 3 times. The polymer solution was then stored at -80 °C and lyophilized. HAGM and HAGM-PEG3-Azido were verified through ^1^H NMR spectroscopy and ATR-FTIR.

#### DBCO and dye conjugation to PAP

Murine recombinant PAP (Abcam) (1 mg/mL) was dissolved in PBS and functionalized with DBCO-PEG4-NHS ester (Vector Labs) in a 1:30 molar excess as per the manufacturer’s recommendation. The protein solution was incubated for 2 h at room temperature, protected from light, and then purified for DBCO-PEG4-PAP using Pierce^TM^ Dye and Biotin Removal Spin columns (Thermo Scientific).

For fluorescent microscopy imaging, PAP was incubated with Alexa Fluor™ 488 NHS Ester (Invitrogen) at a 1:15 molar excess, as recommended by the manufacturer. The labeled PAP was purified using Pierce^TM^ dye removal columns. DBCO-PEG4-PAP was labeled with the above fluorescent compound simultaneously with the reaction with DBCO-PEG4-NHS ester.

#### Fabrication of Cryogel_VAX_ and O_2_-Cryogel_VAX_

Cryogel vaccine scaffolds against prostate cancer were fabricated using HAGM (Cryogel_VAX_) or HAGM-PEG3-Azido (Click Cryogel_VAX_) (4 % w/v) by cryo-polymerization (TEMED, 0.07 % w/v; APS, 0.28 % w/v) at -20 °C. HAGM and HAGM-PEG3-Azido were dissolved in deionized H_2_O with 10 µg PAP and DBCO-PEG4-PAP, respectively, overnight at 4 °C before cryo-polymerization. Immunomodulatory factors (25 µg/cryogel Poly(I:C) and 1.5 µg/cryogel GM-CSF) were incorporated into the polymer solution immediately prior to cryo-polymerization. To produce O_2_-generating cryogel vaccine scaffolds (O_2_-Cryogel_VAX_, Click O_2_-Cryogel_VAX_), cryogels were hybridized with ball-milled calcium peroxide (CaO_2,_ 1 % w/v), and grafted with chemically modified catalase (CAT, 1 % w/v) as previously reported.^57,59^

#### Loading and release of immunomodulatory factors and proteins from cryogels

To determine the loading efficiency and in vitro release kinetics of GM-CSF and PAP from Cryogel_VAX_ and O_2_-Cryogel_VAX_, gel samples were briefly washed in 70 % ethanol followed by 2 PBS washes. Each washed gel was dissociated to measure loading or incubated in sterile PBS in a microcentrifuge tube under orbital shaking at 4 °C. Partial supernatant was removed periodically and replaced with an equal volume of fresh buffer. GM-CSF and PAP were detected by either ELISA (Uncoated mGM-CSF, Invitrogen) or bicinchoninic acid assay (micro-BCA, Thermo Scientific).

#### O_2_ release kinetics (H_2_O_2_ depletion)

O_2_ release of O_2_-Cryogel_VAX_ was monitored using O_2_ optical sensor spots (OXSP5-ADH, PyroScience, GmbH, Aachen, Germany). O_2_-Cryogel_VAX_ scaffolds were individually placed into a 96 well plate containing 200 µL of PBS at room temperature. Dissolved O_2_ (Torr) within the well was recorded for 48 h using FireString software. To ensure adequate H_2_O_2_ depletion through the inclusion of CAT, O_2_-Cryogels were incubated in PBS at 37 °C for 48 h and H_2_O_2_ concentration in the supernatant was measured using the Pierce^TM^ Quantitative Peroxide Assay Kit (Thermo Scientific).

#### Mouse model and study design

Animal experiments were carried out in compliance with the National Institutes of Health (NIH) guidelines and approved by the Division of Laboratory Animal Medicine and Northeastern University Institutional Animal Care and Use Committee (protocol number 23-0512R). Vaccination studies were performed on 6–8-week-old male C57BL/6 mice (Charles River). Bolus_VAX_, O_2_-Cryogels, and Cryogel_VAX_ scaffolds were injected subcutaneously (s.c.) in both flanks close to the inguinal LNs (total of 2 injections/mouse). All cryogel scaffolds were sterilized in 70 % ethanol for 10 min, and washed 3 times with sterile PBS for 5 min before use in animals.

#### Antigen presenting cell characterization in draining LNs

Mice received immunization (D0) and boost (D14) doses of Bolus_VAX_, Cryogel_VAX_, Click Cryogel_VAX_, O_2_-Cryogel_VAX_, Click O_2_-Cryogel_VAX_, or O_2_-Cryogel alone. Blood samples were collected every 7 d. The inguinal LNs were explanted 28 d after initial immunization and dissociated into single-cell suspensions. Cells were stained with fixable viability dye eFluor 506 (eBioscience, 1:400 dilution in PBS) for 30 min at 4 °C. The cells were subsequently washed with protein blocking buffer (PBA) (PBS + 1 % BSA) before being stained overnight with fluorochrome-conjugated antibodies (CD11c-FITC (Clone N418), I-A/I-E-APC-Cy7 (Clone: M5/114.15.2), IgD-Alexa Fluor 700 (Clone 11-26c.2a), CD45-PerCP-Cy5.5 (Clone S18009F), CD11b-PE-Cy7 (Clone: M1/70), F4/80-APC (Clone: BM8), CD19-BV605 (Clone 6D5), BioLegend) in PBA at 4 °C. Cells were washed 3 times with PBA prior to analysis. Flow cytometry measurements were done using the Attune NxT flow cytometer (Thermo Fisher Scientific).

#### Antibody titration by immunoassays

Mouse sera was collected prior to boost immunization on D14 and at endpoint D28. Immunoglobulin isotyping was evaluated using the Mouse Immunoglobulin Isotyping LegendPlex kit (BioLegend) and Uncoated Ig2c Mouse ELISA kit (Thermo Fisher Scientific). Specific IgG to murine PAP was evaluated using a custom ELISA protocol. Specifically, PAP-IgG was detected using recombinant PAP (Abcam) as the capture and anti-IgG (H+L)-biotin (Invitrogen) and streptavidin Poly-HRP (Thermo Scientific) as the detection. Mouse sera were serially diluted to calculate the half-maximal effective concentration (EC50) as the titer for each treatment group.

#### T cell-characterization in spleens

Mice received immunization (D0) and boost (D14) doses of Bolus_VAX_, Cryogel_VAX_, Click Cryogel_VAX_, O_2_-Cryogel_VAX_, Click O_2_-Cryogel_VAX_, or O_2_-Cryogel alone. The spleens were explanted 28 d after initial immunization and dissociated into single-cell suspensions. Cells were stained with fixable viability dye eFluor 506 (eBioscience, 1:400 dilution in PBS) for 30 min at 4 °C. The cells were subsequently washed with PBA before being stained overnight with fluorochrome-conjugated antibodies (CD3ε-FITC (Clone: 500A2), CD44-PE (Clone: IM7), CD4-PerCP-Cy5.5 (Clone: RM4-5), CD8a-APC (Clone: 53-6.7), CD62L-BV605 (Clone MEL-14), BioLegend) in PBA at 4 °C. Cells were washed 3 times with PBA prior to analysis. Flow cytometry measurements were done using the Attune NxT flow cytometer (Thermo Fisher Scientific).

#### Splenocyte activation and intracellular cytokine staining

Splenocytes (40 x 10^6^ cells mL^−1^) were incubated with either Cell Stimulation Cocktail (eBioscience), 30 µg mL^−1^ PAP protein-derived peptides mPAP_114-122_ and mPAP_128-136_ (Biosynth), 0.5 x 10^6^ cells mL^−1^ TRAMP-C2 or control (no stimulation) in Protein Transport Inhibitor Cocktail (eBioscience) solutions for 6 h at 37 °C. Next, the cells were washed with PBS and incubated for 30 min with Fixable Viability Dye eFluor 506 (eBioscience) in PBS (1:400 dilution) at 4 °C. Cells were then washed once with PBS and twice with PBA before being stained for 1 h with fluorochrome-conjugated antibodies (CD3ε-FITC (Clone: 500A2), CD4-APC-Cy7 (Clone: RM4-5), CD8-PerCp-Cy5.5 (Clone: 53-6.7), CD44-BV421 (Clone: IM7), BioLegend) in PBA at 4 °C. Cells were subsequently washed 3 times with PBA, fixed and permeabilized using a FoxP3/Transcription Factor Fixation/Permeabilization Kit (Cell Signaling Technology) according to the manufacturer’s protocol. Intracellular staining was performed by incubating the cells with fluorochrome-conjugated antibodies (IL-2-PE (Clone: JES6-5H4), TNF*α*-AF700 (Clone: MP6-XT22), BioLegend) in permeabilization buffer for 30 min at 4 °C. Lastly, cells were washed 3 times with PBA and analyzed using the NovoCyte Advanteon flow cytometer (Agilent Technologies).

#### Splenocyte activation and IFN-γ ELISpot

Splenocytes (3 x 10^6^ cells mL^−1^) were incubated with either Cell Stimulation Cocktail (eBioscience), 30 ug mL^−1^ PAP protein-derived peptides mPAP_114-122_ and mPAP_128-136_ (Biosynth), 0.5 x 10^6^ cells mL^−1^ TRAMP-C2, or control (no stimulation) in a mouse IFN-γ ELISpot plate (R&D Systems) overnight at 37 °C. Following incubation, the manufacturer’s protocol was followed. Spots were counted manually using a dissection microscope and normalized to the number of cells initially seeded per well.

#### TRAMP-C2 therapeutic model

Male C57BL/6 mice (8–10-week-old) were subcutaneously injected with 2.5 x 10^6^ TRAMP-C2 cells mixed 1:1 with Matrigel Matrix Basement Membrane HC (Corning, 18.97 MG/ML). Mice were immunized with Bolus_VAX_, O_2_-Cryogels, O_2_-Cryogel_VAX_, or Click O_2_-Cryogel_VAX_ when tumors reached a mean volume of 50 mm^3^. Mice were boosted 14 days later, and tumor volume was measured using calipers and recorded two or three times per week until tumors reached the euthanasia criteria endpoint (mean volume >1500 mm^3^). To assess the antitumor immune response, tumors were explanted and dissociated, and infiltrating T cells were analyzed for immunosuppressive markers through flow cytometry using the NovoCyte Advanteon (Agilent Technologies). Cells were stained for 1 h with fluorochrome-conjugated antibodies (CD4-PerCP/Cy5.5 (Clone: RM4-5), CD8-APC (Clone: 53-6.7), CD25-BV785 (Clone: PC61), and CD279 (PD-1)-BV510 (Clone: 29F.1A12), BioLegend) in PBA at 4 °C. Cells were subsequently washed 3 times with PBA, fixed and permeabilized using a FoxP3/Transcription Factor Fixation/Permeabilization Kit (Cell Signaling Technology) according to the manufacturer’s protocol. Intracellular staining was performed by incubating the cells with fluorochrome-conjugated antibody (FOXP3-PE (Clone: MF-14), BioLegend) in permeabilization buffer for 30 min at 4 °C. Lastly, cells were washed 3 times with PBA and analyzed.

### Statistical analysis

Flow cytometry data were processed using FlowJo software and manual gating was performed. Statistical analysis was conducted using GraphPad Prism 10 software (La Jolla, CA, USA). Significant differences between groups were assessed using one-way analysis of variance (ANOVA), mixed-effects analysis, or log-rank (Mantel-Cox) with Bonferroni post hoc tests. Differences were considered statistically significant at *p < 0.05, **p < 0.01, ***p < 0.001, and ****p < 0.0001.

## Acknowledgements

S.A.B. and S.M.H. gratefully acknowledge the financial support from the National Institutes of Health (NIH, 1R01EB027705). S.A.B. also acknowledges funding from the National Science Foundation (NSF CAREER, DMR 1847843) and the “*Chaire d’Excellence de Normandie*”. The authors thank the Institute for Chemical Imaging of Living Systems (CILS) at Northeastern University for imaging support and access to the Zeiss LSM 800 microscope.

## Authorship contribution

Conceptualization, A.N., K.B., T.C., S.A.B. and S.M.H.; methodology, A.N., K.B., T.C., S.A.B. and S.M.H.; investigation, A.N., E.T., L.W., B.N., E.C., L.L.M.; data curation, A.N.; formal analysis, A.N.; supervision, M.S., S.A.B. and S.M.H.; writing—original draft, A.N.; writing—review & editing, A.N., S.A.B, and S.M.H.; funding acquisition, S.A.B. and S.M.H. All authors have reviewed the manuscript and approved its contents.

## Supplementary information

