## Supplemental Information for "Oxygen-generating cryogel vaccines help overcome tumor antigen tolerance and induce durable anti-tumor immunity in prostate cancer"

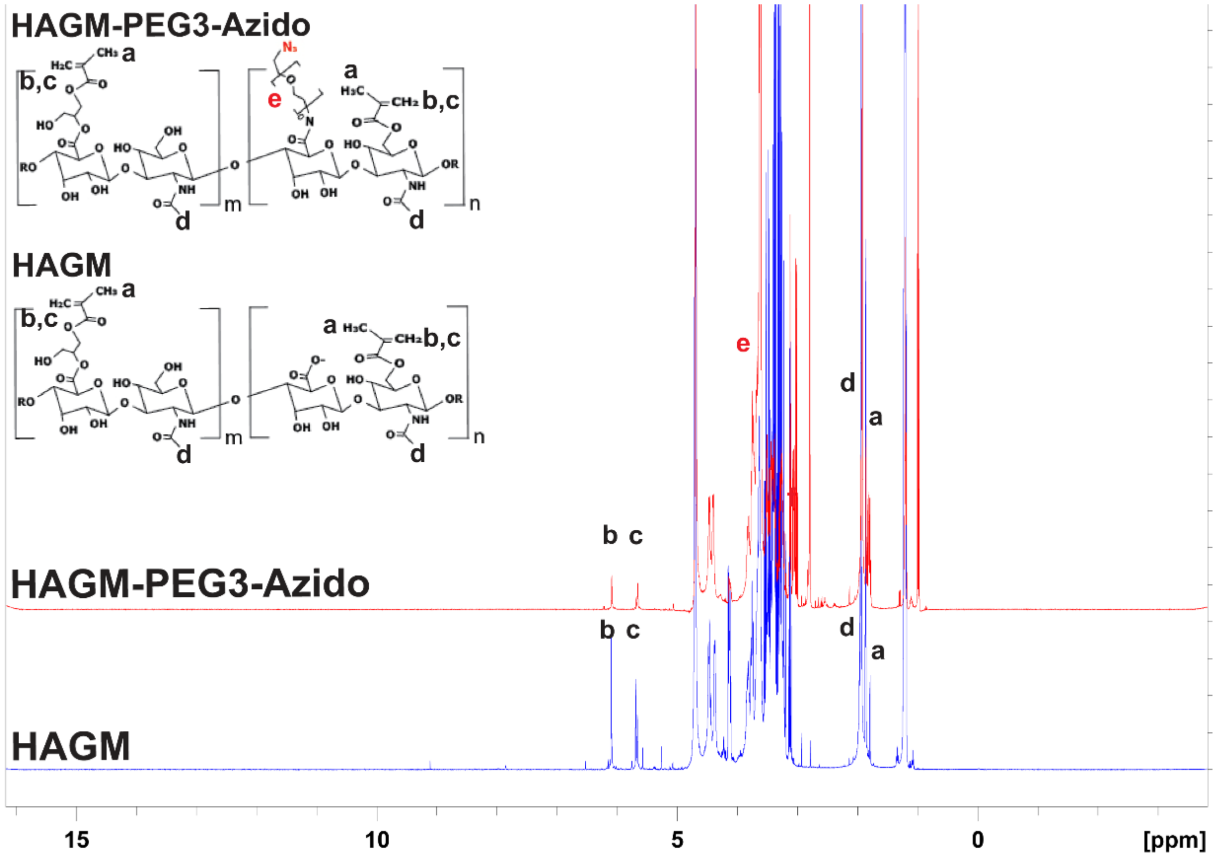


**Figure S1. $^1$H NMR characterization of HAGM and HAGM-PEG3-Azido.** $^1$H NMR spectra of HAGM and HAGM-PEG3-Azido in $D_2O$ with corresponding peaks of methacryloyl and PEG residues.

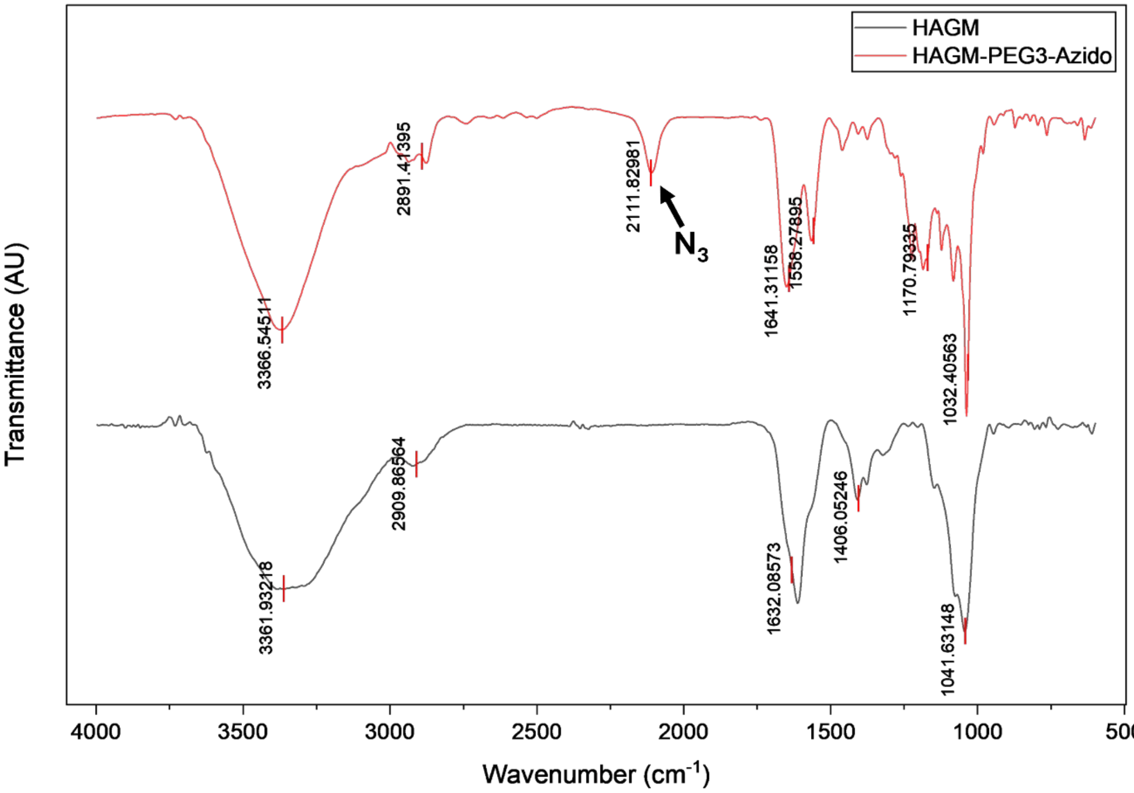


**Figure S2. ATR-FTIR characterization of HAGM and HAGM-PEG3-Azido.** ATR-FTIR spectra of HAGM and HAGM-PEG3-Azido with the introduction of a characteristic $N_3$ peak at 2111 $cm^{-1}$ corresponding to the presence of the azido group in the modified polymer.

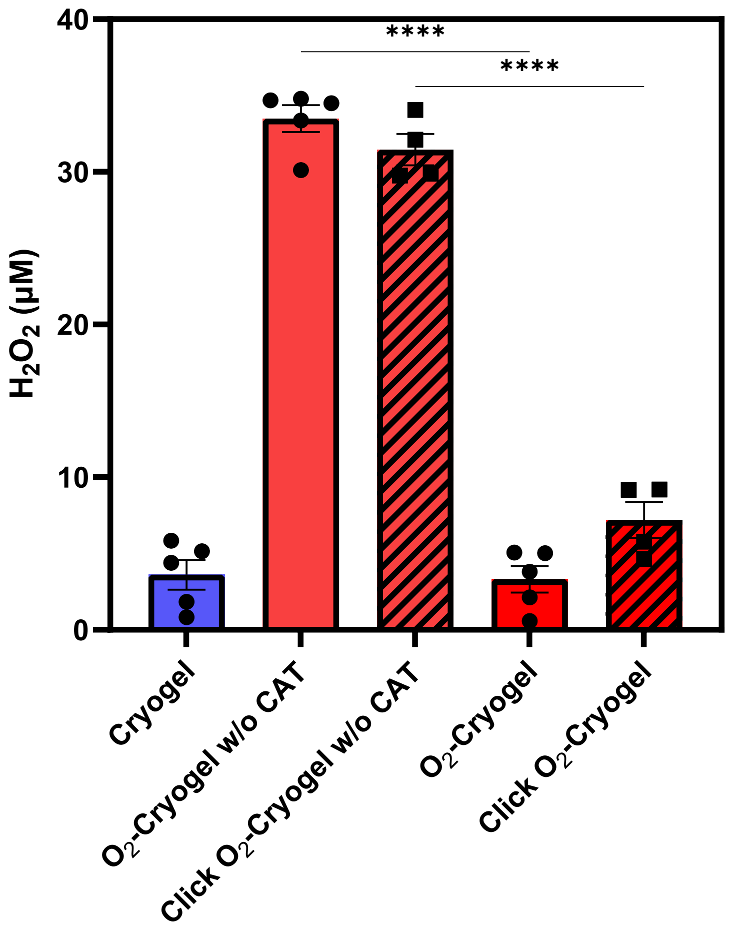


**Figure S3. $H_2O_2$ release from cryogels.** $H_2O_2$ release from $O_2$-Cryogels and Click $O_2$-Cryogels after incubation in PBS for 48 h at 37 °C, formulated without and with catalase (CAT, 1% w/v), with non-oxygen-releasing cryogels as a negative control. Data are presented as mean ± SEM (n = 4–5). Statistical analysis was performed using one-way ANOVA and Tukey's post hoc test: ****$p < 0.0001$.

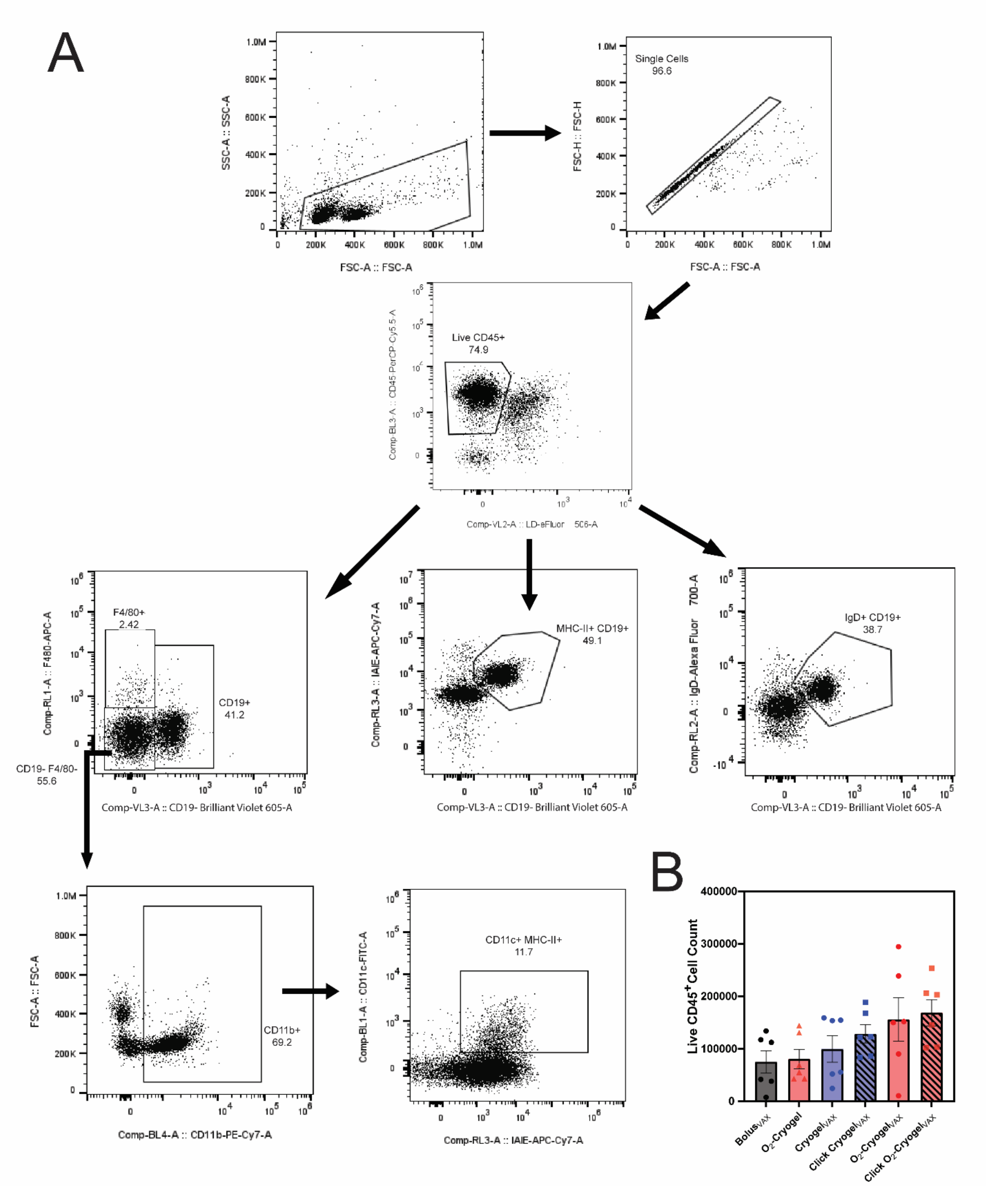


**Figure S4. Analysis of APCs in inguinal lymph nodes (LNs) by flow cytometry. (A)** Gating strategy for dendritic cells ($CD11b^+$, $CD11c^+$, IA-$IE^+$), macrophages ($F4/80^+$), antigen-presenting (IA-$IE^+$, $CD19^+$) and mature B cells ($IgD^+$, $CD19^+$) collected from the draining LNs 28 d after initial vaccination. **(B)** Cell count of live $CD45^+$ cells in inguinal LNs.

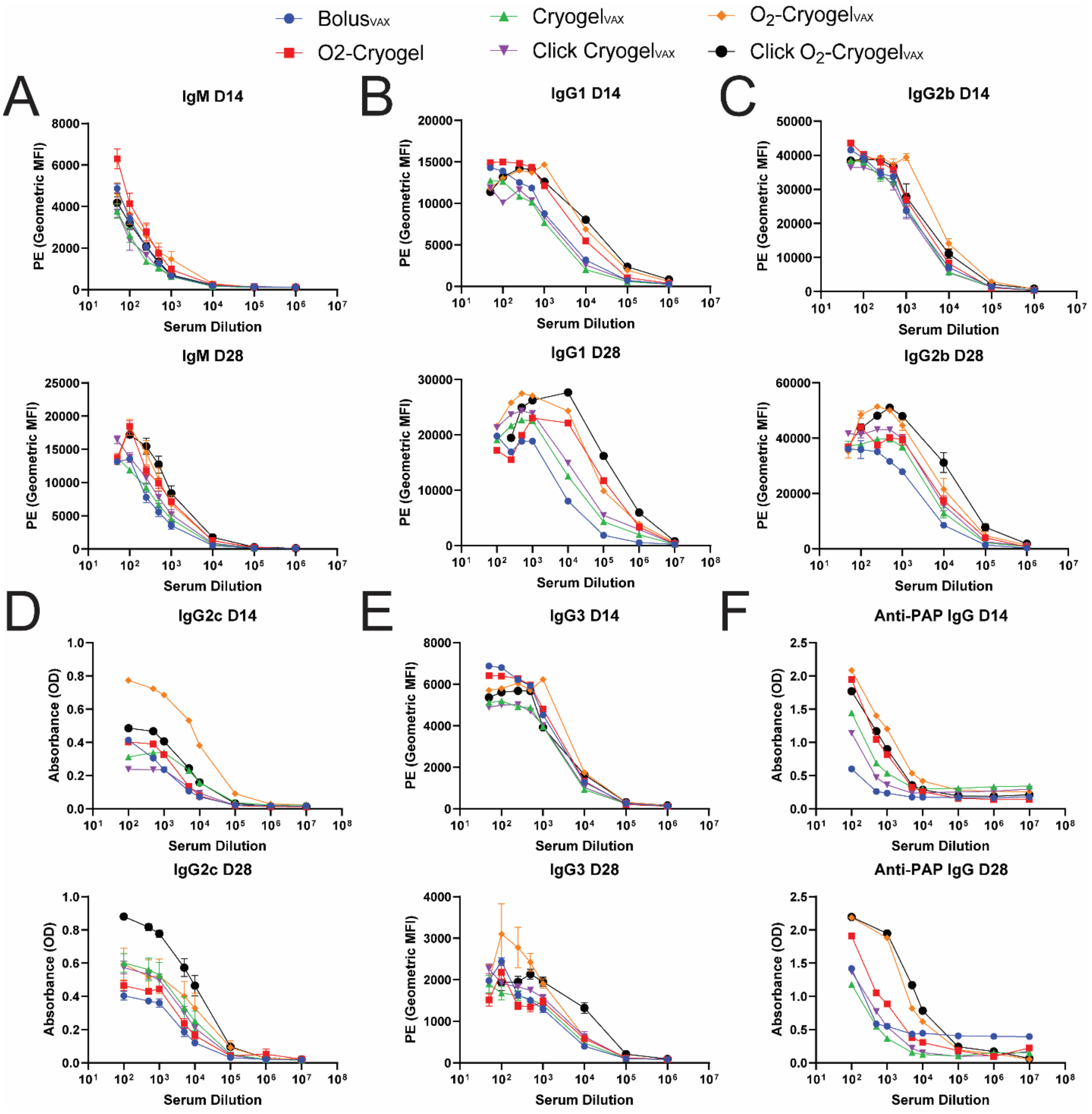


**Figure S5. Serological antibody assay titer curves.** Mouse sera levels for **(A)** IgM, **(B)** IgG1, **(C)** IgG2b, **(D)** IgG2c, **(E)** IgG3, and **(F)** PAP-specific IgG 14 d after priming (D14) and boosting (D28) with various vaccine formulations.

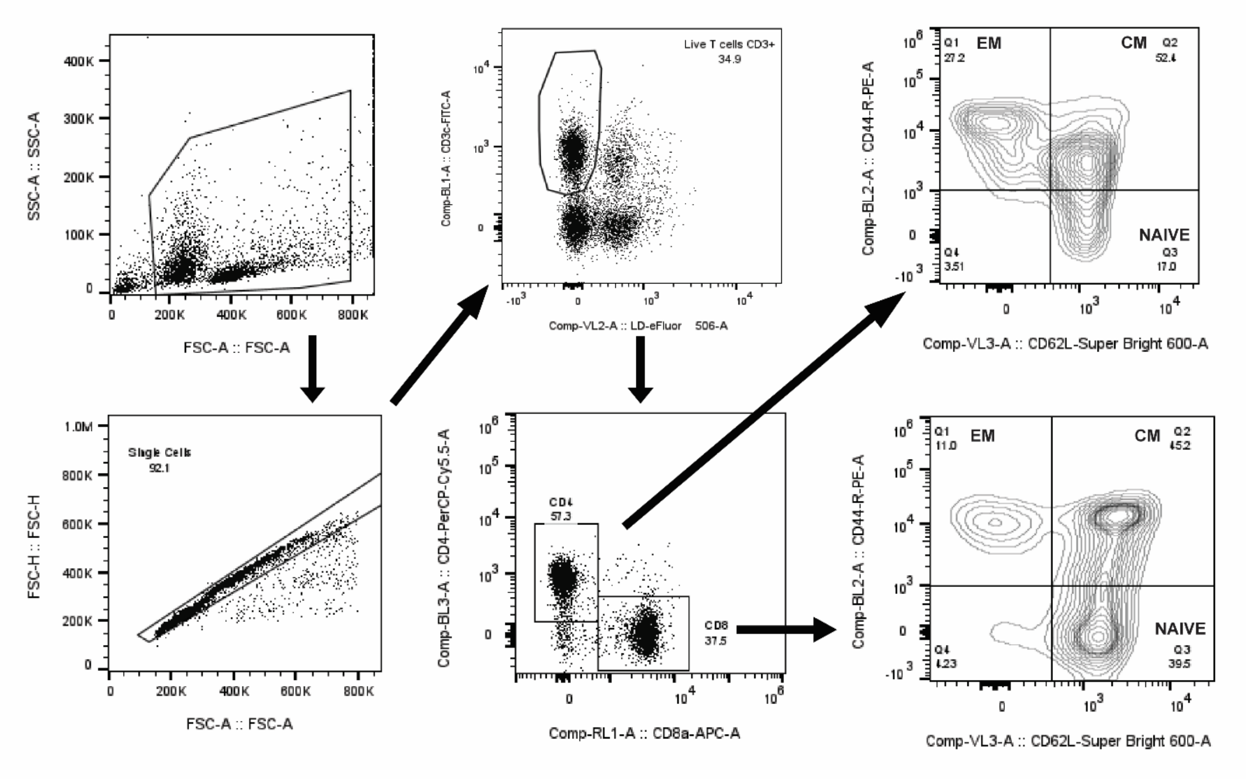

**Figure S6. Analysis of memory T-cell populations in spleens by flow cytometry**. Gating strategy for $CD4^+$ and $CD8^+$ effector memory (EM) ($CD44^+$, $CD62L^-$), central memory (CM) ($CD44^+$, $CD62L^+$), and Naïve ($CD44^-$, $CD62L^+$) T cells collected from the spleens 28 d after initial vaccination.

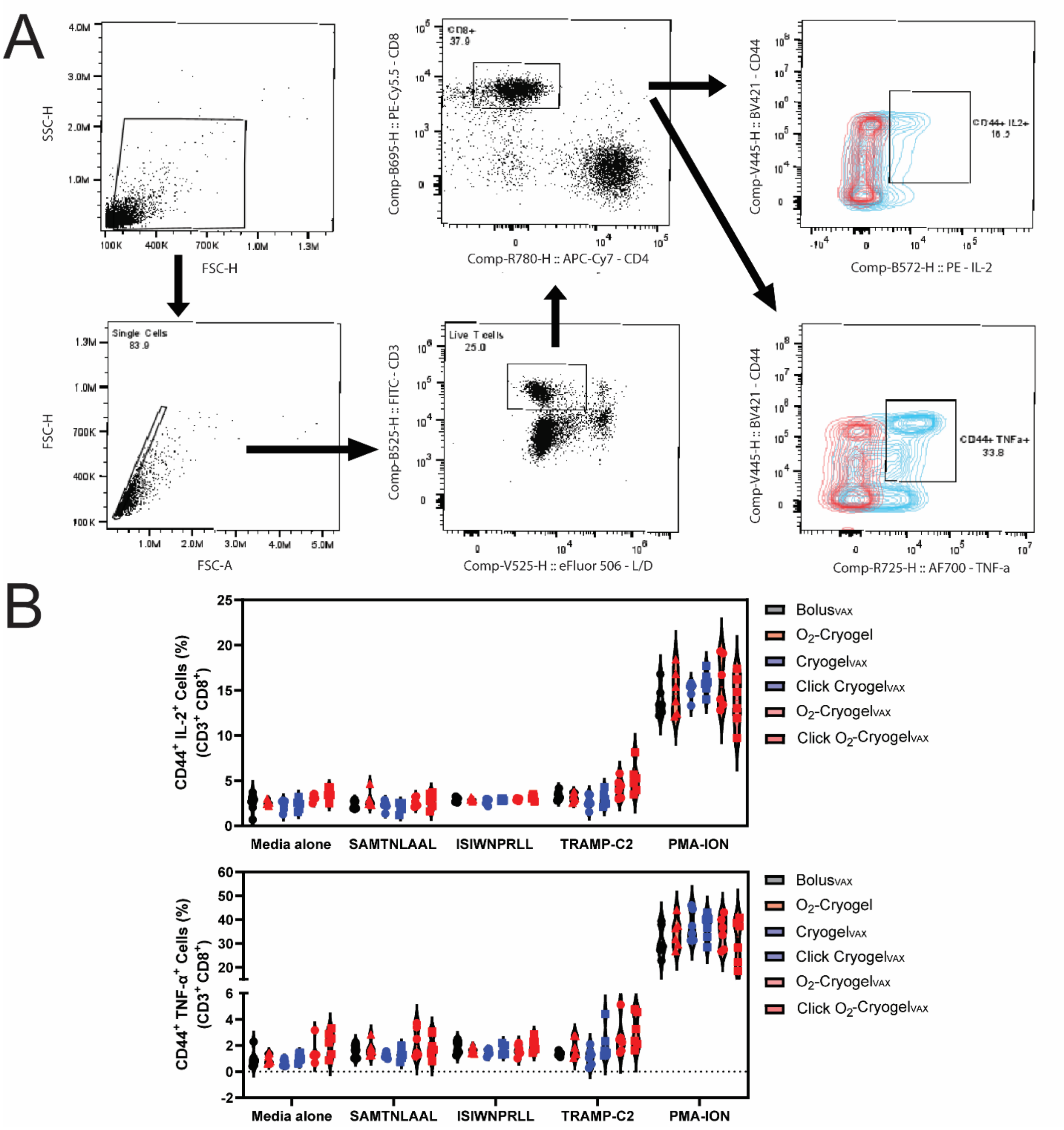


**Figure S7. Splenocyte activation analysis by flow cytometry. (A)** Gating strategy for $CD8^+$ T-cell intracellular cytokines ($IL\text{-}2^+$, $TNF\text{-}\alpha^+$) after co-incubation of collected splenocytes with stimuli for 6 h at 37 °C. **(B)** Percentages of $CD44^+$, $IL\text{-}2^+$ and $TNF\text{-}\alpha^+$ cells after incubation with either media alone, PAP-derived peptides: $mPAP_{114\text{-}122}$ and $mPAP_{128\text{-}136}$,TRAMP-C2 cells, or PMA-Ionomycin (PMA-ION) within splenocytes collected from vaccinated mice with various formulations.

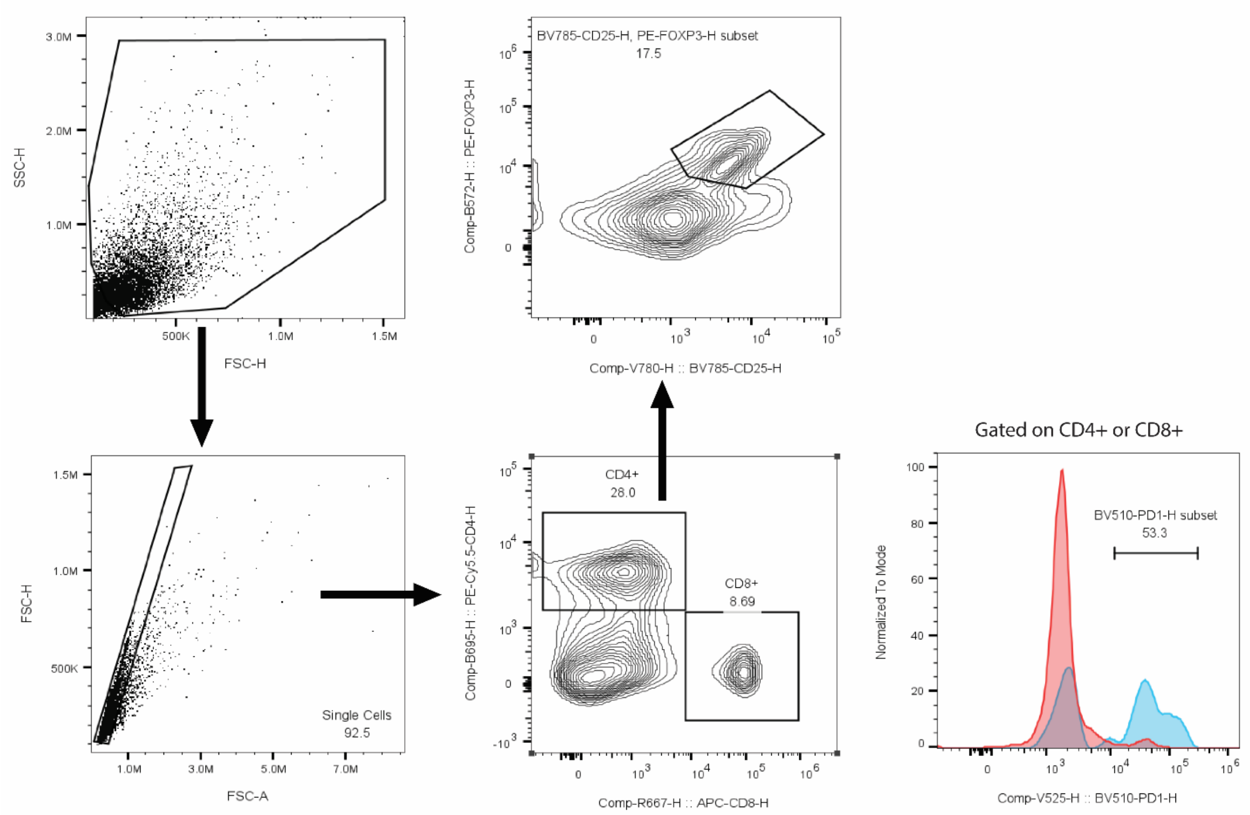


**Figure S8. Analysis of tumor-infiltrating T cells by flow cytometry.** Gating strategy of regulatory T cells ($CD4^+$, $CD25^+$, $FoxP3^+$) and PD-1-expressing T cells ($CD4^+/CD8^+$, $PD\text{-}1^+$) collected from TRAMP-C2 subcutaneous tumors.
